# Rapamycin enhances VEGFA and VSMC contractile protein expression in MSCs via mTORC1 inhibition

**DOI:** 10.1101/2025.08.05.668760

**Authors:** Pedro Bartkevitch Rodrigues, Sai Sanikommu, Omar Elwardany, Tiffany A. Eatz, Rachele Angiolini, Helena Hernandez-Cuervo, Naser Hamad, Jennifer Sophirum Suon, Maxon V. Knott, Joshua M. Hare, John W. Thompson, Robert M. Starke

**Affiliations:** Department of Neurological Surgery, Radiology and the University of Miami Cerebrovascular Initiative, University of Miami; Neurosurgery Department, Faculty of Medicine, Suez Canal University, Ismailai, Egypt; The Interdisciplinary Stem Cell Institute, Miller School of Medicine, University of Miami, Miami, FL, USA

**Keywords:** mTOR, Rapamycin, MSCs, VSMC markers, Angiogenesis, VEGFA, Cerebral aneurysms

## Abstract

**Introduction:** Intracranial aneurysm (ICA) rupture is the most common cause of non-traumatic subarachnoid hemorrhage, a devastating type of stroke. Vascular smooth muscle (VSMC) and endothelial cell (EC) dysfunction and death play important roles in the etiology of ICA formation, rupture and treatment failure. Mesenchymal stem cells (MSCs) have been extensively investigated for their therapeutic potential in vascular diseases. The mammalian target of rapamycin (mTOR) is a key regulatory pathway involved in cellular functions controlling intracellular anabolic and regulatory processes. Rapamycin, a specific mTOR complex 1 inhibitor, is widely used in the clinical management of cardiac and vascular pathologies. In this study, we explored MSC’s potential to express VSMC and EC markers under the influence of rapamycin.

**Methods:** Human MSCs were treated with rapamycin for 2, 5, and 10 days, and cell death and proliferation determined by MTT analysis. Protein expression and phosphorylation levels were determined by western blot analysis. Calcein AM and PI staining was used to determine cell viability, and morphology.

**Results:** MTT and Calcein AM analysis demonstrated that prolonged rapamycin treatment did not affect MSC viability but was found to reduce MSC proliferation and cause a nearly threefold increase in cell size compared to controls. Rapamycin treatment increased expression of the VSMC protein markers, α-SMA, and transgelin and was required to maintain elevated expression levels both proteins. In contrast, selective inhibition of mTORC2 with JR-AB2-01 caused a decrease in α-SMA and trangelin expression, suggesting mTORC1 regulation of α-SMA and transgelin expression, rather than mTORC2. Rapamycin treated MSC were also found to induce cell sprouting between adjacent cells. MSCs were found not to express endothelial protein markers under either basal conditions or following rapamycin treatment. However, rapamycin treatment was found to increase VEGFA expression, a known promoter of angiogenesis.

**Conclusion:** Rapamycin has the potential to increase VSMC protein expression in MSCs by its action on mTORC1. The ability of rapamycin to increase VEGFA expression in MSCs could allow for a novel MSC-based therapeutic intervention in cerebral aneurysm management which could aid in ICA healing and reducing rupture risk.

## 1. INTRODUCTION

Intracranial aneurysms (ICA) are a dilation or bulging of a cerebral artery which is present in approximately 3.2% of the general population (1). The ICA can become unstable and rupture, resulting in a subarachnoid hemorrhage, a type of stroke associated with high rates of morbidity and mortality (2). Treatment options for ICAs are only mechanical and include the placement of metal clips across the aneurysm neck or the endovascular placement of occlusion devices, such as coils within the aneurysm or stents across the aneurysm orifice. Since endovascular intervention is less invasive and associated with better outcomes, it has become the first-line treatment modality for ICA. However, endovascular treatment is associated with aneurysm recanalization and treatment failure, which can occur due to the slow and incomplete reformation of the parent artery across the aneurysm neck.

Vascular smooth muscle (VSMC) and endothelial (EC) cell dysfunction is a hallmark of aneurysmal formation and rupture (3–5). It is believed that EC dysfunction occurs early in ICA development and initiates an inflammatory cascade characterized by inflammatory cell infiltration, inflammatory cytokine expression, and extracellular matrix degradation. As a result, VSMCs undergo a phenotypic switch from a contractile to a synthetic phenotype which further exacerbates tissue inflammation by the expression of inflammatory cytokines and matrix degrading protein (6–11). As the aneurysm develops changes in hemodynamic stress causes further VSMC dysfunction and apoptosis, vessel wall thinning and ultimately, ICA rupture (12–18).

Mesenchymal stem cells (MSCs) are stromal-derived cells capable of self-renewal and differentiation into various cell types, including osteoblasts, chondrocytes, myocytes, and adipocytes (19–23). As such, MSCs have been extensively investigated as a potential cell therapy for many diseases, including myocardial infarction (24–27), pulmonary fibrosis (28), systemic lupus erythematosus (29), and others (30–33). In ICA, MSC coated endovascular coils were found to improve aneurysm healing and endothelialization of the aneurysm neck in a rabbit aneurysm model (34). However, the therapeutic potential of MSC for ICA treatment has not been fully investigated.

Mammalian target of rapamycin (mTOR) is a serine/threonine kinase, which functions through forming two distinct multiprotein complexes: mTOR Complex 1 (mTORC1) and Complex 2 (mTORC2) (35). mTOR serves as a “hub” integrating extracellular and intracellular signals into changes in cellular activity. As such mTOR is a known regulator of the cell cycle, lipid and glucose metabolism, and nucleotide and protein synthesis (35). Rapamycin, a mTORC1 specific inhibitor, is used in the treatment of restenosis in cardiac and vascular pathologies by reducing VSMC proliferation (36–40). In preclinical studies, rapamycin promotes human SMC differentiation (41) and embryonic stem cells differentiation into cardiomyocytes in mice (42). However, the potential role of rapamycin in driving MSCs differentiation into VSMCs remains unexplored and warrants further investigation.

In this study, we explored the potential of rapamycin to induce MSCs expression of VSMC and EC genes which may allow the therapeutic use of MSCs in ICA parent vessel healing.

## 2. MATERIALS AND METHODS

### 2.1 Cell culture

Umbilical cord-derived MSCs CD105, CD73, CD90, CD45, CD34, CD14, CD19, HLA-DR; Sigma-Aldrich) were cultured in a growth medium consisting of alpha Minimum Essential Medium (αMEM) supplemented with 10% fetal bovine serum (FBS) and 1% penicillin/streptomycin/glutamine (P/S/G). Passages 3-10 were used for the experiments. For experiments, MSCs were seeded at a density of 5 × 10^^5^ cells per 100 mm dish and allowed to adhere overnight. Cells were treated with rapamycin (ThermoFisher) or the specific mTORC2 inhibitor JR-AB2-01 (Sigma) at various concentrations and times. Cells treated with vehicle (DMSO) served as controls. For long-term rapamycin exposure (> 72hrs), cells were fed fresh media containing rapamycin or vehicle (for control cells).

### 2.2 MTT (viability, cytotoxicity and proliferation analysis)

Cell viability and proliferation were assessed using MTT (3-(4,5-dimethylthiazol-2-yl)-2,5-diphenyltetrazolium bromide) assay (Sigma). Cells were treated with increasing concentrations of rapamycin (0.005, 0.01, 0.05, 0.1, 1, 10, and 1000 ng/mL) for 48 and 72 hours. After treatment, 100 µL of MTT solution (0.5 mg/mL) was added to each well of a 96-well plate and incubated for 4 hours. The medium was carefully removed, and 100 µL of DMSO was added to dissolve the formazan crystals. MTT reaction was measured by absorbance at 570 nm. Cell proliferation was expressed as relative absorbance normalized to untreated controls.

### 2.6 Microscopy Imaging for Assessing Viability, Proliferation, and Morphological Changes

MSCs were treated with rapamycin (10 ng/mL and 500 ng/mL) or vehicle control for 1, 5, 7, 14, and 21 days and cellular proliferation was determined by brightfield microscopy. To assess viability, MSCs were treated with rapamycin (0.1, 1, and 10 ng/mL) for 72 hours and then incubated with Calcein AM (2µM; Invitrogen) to label viable cells and propidium iodide (PI; Fluka) to label non-viable cells. Quantitative morphological analysis was performed by calculating the length-to-width ratio and area of individual cells to determine changes in cell morphology using ImageJ software.

### 2.4 Western blotting

Cells were lysed in ice-cold RIPA buffer containing protease and phosphatase inhibitors (Thermoscientific, Pierce). Protein concentration was measured using BCA assay. Equal amounts of protein (20–30 µg) were separated on 10% to 15% SDS-polyacrylamide gels and transferred onto nitrocellulose membrane. Membranes were blocked with 5% nonfat milk in TBS-T (25 mM Tris, 137 mM NaCl, 2.7 mM KCl, 0.05% Tween-20) for 1 hour at room temperature. The membranes were incubated overnight at 4°C with the primary antibodies: α-SMA (Abcam), Transgelin (Santa Cruz Biotechnology), S6 (Cell Signaling Technology), p-S6 (Cell Signaling Technology), AKT (Cell Signaling Technology), p-AKT (Cell Signaling Technology), Vascular Endothelial Growth Factor A (VEGFA) (Abcam), α-tubulin (CALBIOCHEM). The membranes were washed and incubated with HRP-conjugated secondary antibodies (Bio–Rad) for 1 hour at room temperature. Protein bands were visualized using enhanced chemiluminescence reagents and captured using a chemiluminescence imaging system. Band intensities were quantified using ImageJ software and normalized to α-tubulin.

### 2.7 Statistical analysis

All data were expressed as mean ± standard error of the mean (SEM). Statistical analysis between two groups was performed using the unpaired Student’s t-test. Statistical analysis between more than two groups was performed using a one-way ANOVA with Dunnett’s multiple comparison post hoc test. P < 0.05 was considered statistically significant.

## 3. RESULTS

### 3.1 Rapamycin reduces MSC proliferation

Rapamycin is a known inhibitor of cellular proliferation. Therefore, to better understand the effects of rapamycin on MSCs, we assessed MSC proliferation following treatment with increasing concentrations of rapamycin (0.005, 0.01, 0.05, 0.1, 1, 10, and 1000 ng/mL) for 48 and 72 hours using an MTT assay. As shown in Figure 1a, there was a concentration-dependent decrease in MSC proliferation 48 hours following rapamycin treatment. The lowest rapamycin concentrations (0.005 and 0.01 ng/mL) did not affect MSC proliferation; however, there was a gradual, significant decrease in MSC proliferation, reaching 65% of the control at the highest concentration of rapamycin used. Similarly, rapamycin treatment for 72 hours induced a significant concentration-dependent reduction in cell proliferation, with each concentration reaching 30% of the control at the highest rapamycin concentration used. These results were confirmed using bright-field microscopy. It was found that control MSC cultures reached near confluency within 5 days. In contrast rapamycin-treated MSCs were still below 50% confluency even at 1 week. (Figure 1b).

**Figure 1.**
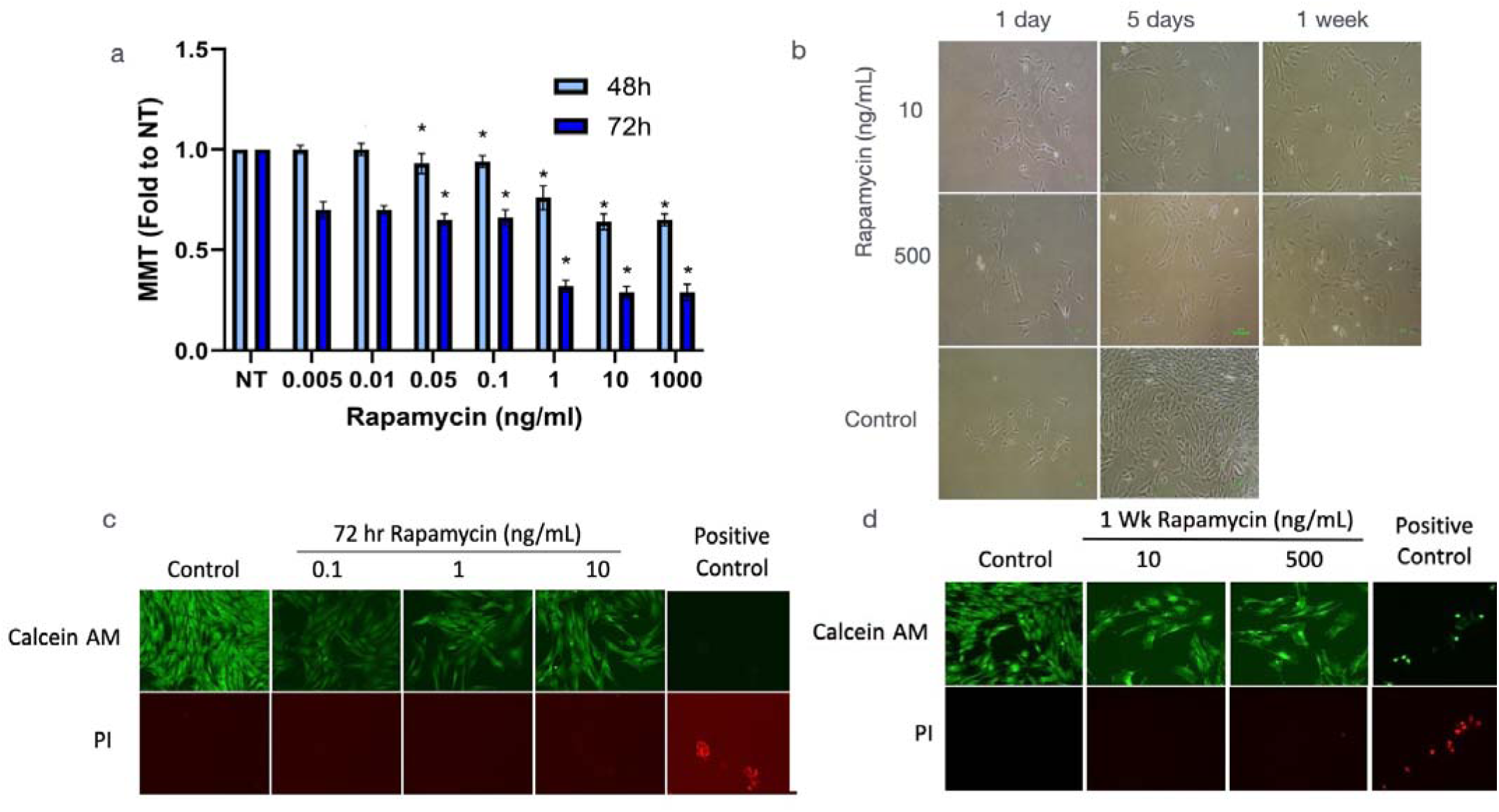
Effects of rapamycin on MSC proliferation and viability. (a) Dose- and time-dependent effects of rapamycin on MSC proliferation measured by MTT assay. Data represent fold change in MTT activity compared to vehicle (NT). In (b) bright-field microscopy illustrating MSC morphology over a 3 week in the presence of 10ng/ml and 500ng/ml rapamycin treatment vs NT. MSC viability was determined by Calcein AM and PI staining after 72 hours (c) and (d) 1 week rapamycin treatment. As a positive control, cells were treated with 0.6% triton x-100 for 10 min. Data are represented as mean ± SEM bars. N = 10, \**P* < 0.05, compared to NT.

### 3.2 Rapamycin does not affect MSC viability

To confirm that rapamycin was affecting cell proliferation and not viability, we then evaluated the cytotoxic effects of rapamycin using the viability dyes, Calcein AM and propidium iodide (PI). MSCs were treated with rapamycin at 0.1, 1, and 10 ng/mL for 72 hours, and cell viability determined at 72hrs and 1 week. As shown in Figure 1c, strong Calcein AM staining was observed in untreated control cells, which was similarly observed in rapamycin-treated cells. PI staining was not observed in any of the cultures treated with rapamycin but was observed in cells treated with 0.6% triton X-100 for 10min (positive control). Consistent with our MTT results, there was a clear reduction in cell numbers in rapamycin-treated cultures (Figure 1c). To assess whether higher concentrations and longer exposures of rapamycin induce cytotoxicity, we treated MSCs with 10 ng/mL or 500 ng/mL rapamycin for one week, and Calcein AM and PI staining determined viability. As shown in Figure 1d, untreated control cells remained viable with strong Calcein AM staining and no PI staining. At 10 ng/mL and 500 ng/mL rapamycin treatment, there was a clear reduction in cell density compared to controls and strong Calcein AM staining with no PI staining except in positive control treated cells. These findings demonstrate that rapamycin inhibits MSC proliferation but is not cytotoxic even at high rapamycin concentrations and prolonged rapamycin exposure.

### 3.3 Rapamycin induced morphological changes and increased cell area

During the analysis of MSC proliferation and viability, it was observed that MSCs underwent changes in cellular morphology in the presence of rapamycin. To further characterize these alterations, fluorescence microscopy was used to assess both cell area and length-to-width ratio. Consistent with other studies, under normal conditions (43), MSCs displayed a typical spindle-shaped morphology (Figure 2a). However, MSCs treated with rapamycin for 72 hours demonstrated a broader and more polygonal shape (Figure 2a). Quantification of cell area revealed a significant increase following rapamycin treatment, with treated cells exhibiting nearly a threefold increase in area compared to controls (*p < 0.05*, Figure 2b). Additionally, cell shape was characterized by calculating the length-to-width ratio. In untreated controls, cells exhibited a ratio of 6.07 (± 1.85), consistent with an elongated morphology. However, in rapamycin-treated cells, the length-to-width ratio was significantly reduced (2.47 ± 0.98), indicating a transition to a more cuboidal cellular profile (*p < 0.05*, Figure 2c). These results demonstrate that rapamycin induces changes in MSC physiology that extend beyond regulating proliferation.

**Figure 2:**
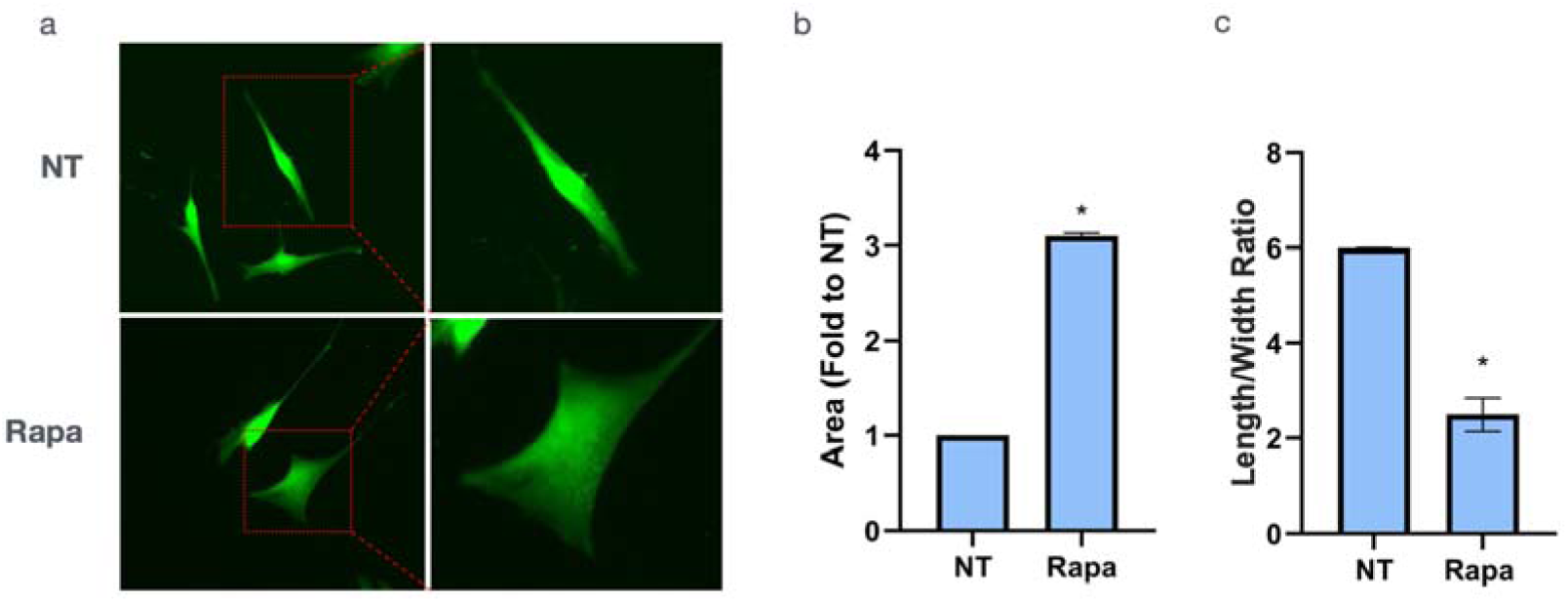
Rapamycin induces morphological changes and area quantification of MSCs following rapamycin treatment. (a) Bright-field microscopy images of control and rapamycin-treated MSCs (10ng/ml) for 72 hours. (b) Quantification of cell area relative to no-treatment control, showing a significant 3-fold increase in Rapamycin-treated cells. (c) Quantification of MSC length-to-width ratio following rapamycin treatment. Data are represented as mean ± SEM bars. N = 10, \**P* < 0.05, compared to NT

### 3.5 Rapamycin promoted vascular smooth muscle cell marker expression in MSCs

The long-term goal of this research is to better understand the therapeutic potential of MSCs in vessel healing following ICA treatment. Based on this goal and the observed morphological changes induced by rapamycin, we next investigated whether rapamycin could stimulate the expression of VSMC markers in MSCs. To test this, MSCs were exposed to 10 ng/mL rapamycin, and the protein expression levels of α-SMA and transgelin were determined following 2, 5, and 10 days of rapamycin treatment. As shown in Figure 3a, under normal conditions, MSCs express a basal level of α-SMA and transgelin. Treating cells with rapamycin resulted in a progressive increase in α-SMA expression over time. At 2 days, α-SMA levels were elevated 1.9-fold relative to controls (*p < 0.05*), increasing further to 2.2-fold at 5 days (*p < 0.05*), and reaching 2.5-fold at 10 days (*p < 0.05*) (Figure 3b). Similarly, transgelin expression was upregulated at all time points examined. Compared to controls, transgelin expression was significantly increased 1.3-fold at 2 days (*p < 0.05*), 1.4-fold at 5 days (*p < 0.05*), and 1.9-fold at 10 days (*p < 0.05*) (Figure 3c). These results demonstrate that MSC expression of VSMC contractile protein expression can be upregulated by rapamycin.

**Figure 3:**
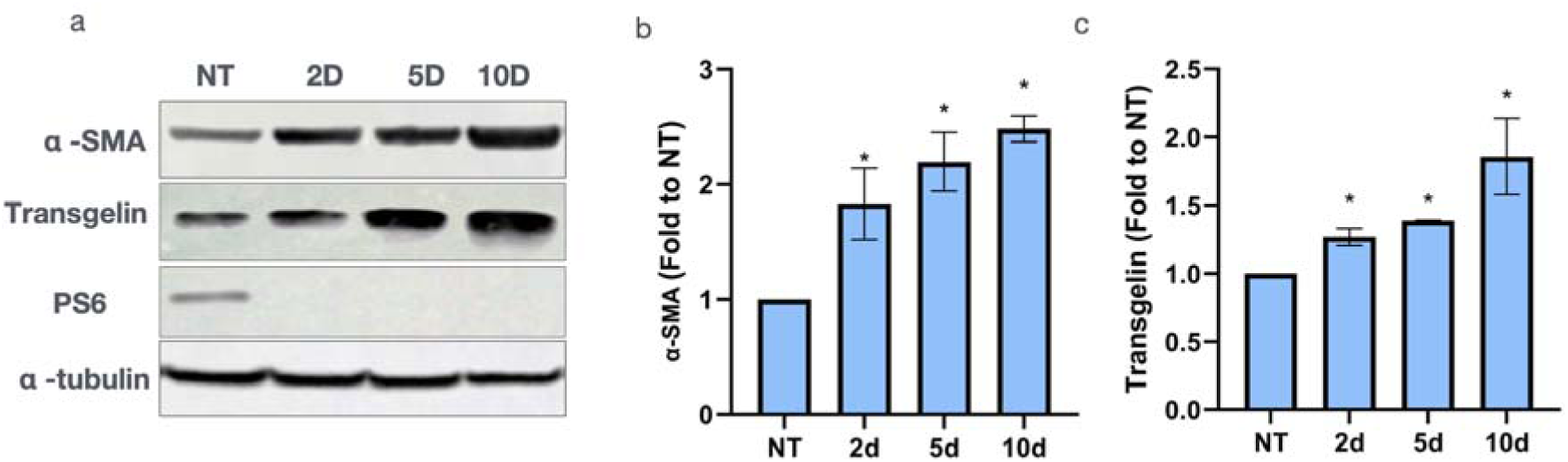
Time-dependent upregulation of VSMC proteins by rapamycin. (a) Western blot analysis showing the expression of α-SMA, transgelin, and α-tubulin in MSCs treated with rapamycin 10ng/ml for 2, 5, and 10 days. Western blot quantification is shown in (b and c). Data are represented as mean ± SEM bars. N = 10, \**P* < 0.05, compared to NT.

### 3.6 Rapamycin Withdrawal Reduces VSMC Marker Expression in MSCs

To determine whether continued rapamycin exposure is required to maintain VSMC marker expression, MSCs were treated with rapamycin (10ng/ml) for 2 days to induce α-SMA and transgelin expression (as shown in Figure 3), followed by either continued rapamycin treatment or drug withdrawal for an additional 3 days. Transgelin and α-SMA protein levels were then assessed and compared to those of untreated controls. As expected, continuous rapamycin exposure resulted in a 2.7-fold increase (*p < 0.05*) in α-SMA and a 1.9-fold increase (*p < 0.05*) in transgelin expression when compared to controls. In contrast, α-SMA and transgelin protein expression were significantly reduced by rapamycin withdrawal. Rapamycin withdrawal reduced α-SMA expression compared to continuous-rapamycin-treated cultures, but it was still significantly increased, 1.4-fold (*p < 0.05*), compared to control, untreated MSC (Figure 4b). Likewise, transgelin protein expression returned to control baseline levels following rapamycin withdrawal (Figure 4c).

**Figure 4:**
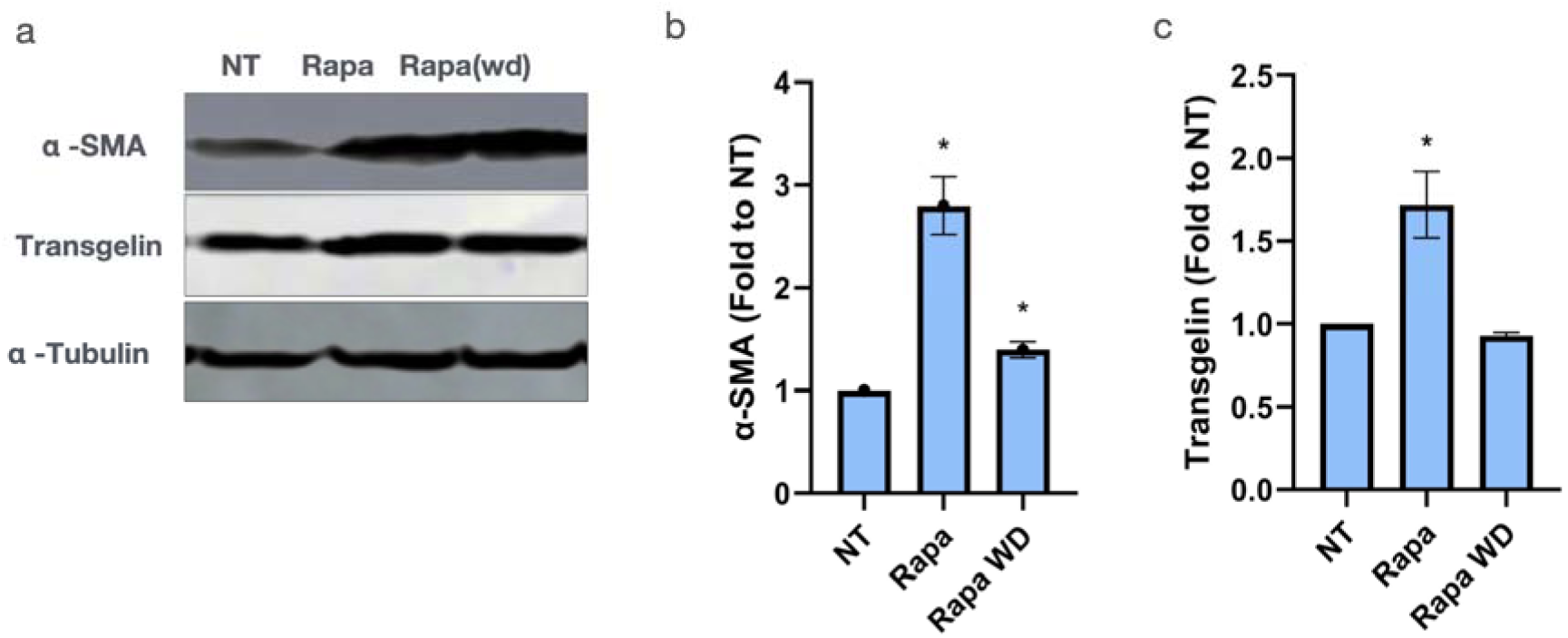
Rapamycin withdrawal reduces VSMC marker expression in MSCs. a) α-SMA, and transgelin protein expression following rapamycin treatment for 7 days, and rapamycin treatment for 2 days followed by 5 days without rapamycin (Rapa wd). Western blot quantification is shown in (b) and (c). Data are represented as mean ± SEM bars. N = 10, \**P* < 0.05, compared to NT.

### 3.7 Rapamycin Suppresses mTORC2 Activity in MSCs

Rapamycin is a well-established specific inhibitor of mTORC1. However, rapamycin can indirectly inhibit mTORC2 during prolonged exposure to rapamycin (44). Therefore, we were interested in determining whether rapamycin inhibited mTORC2 activity in our MSCs. To test this, cells were treated with rapamycin (10 ng/mL) for 2, 5, and 10 days, and mTORC2 activity was determined by changes in the phosphorylation of a downstream mTORC2 target protein, AKT (Ser473). As shown in Figure 5a and b, rapamycin treatment reduced AKT (Ser473) phosphorylation by 12% at 2 days of treatment compared to untreated controls (*p < 0.05*), and was further decreased with time following rapamycin exposure, declining by 18% at 5 days (*p < 0.05*) and 20% at 10 days (*p < 0.05*) (Figure 5b).

**Figure 5:**
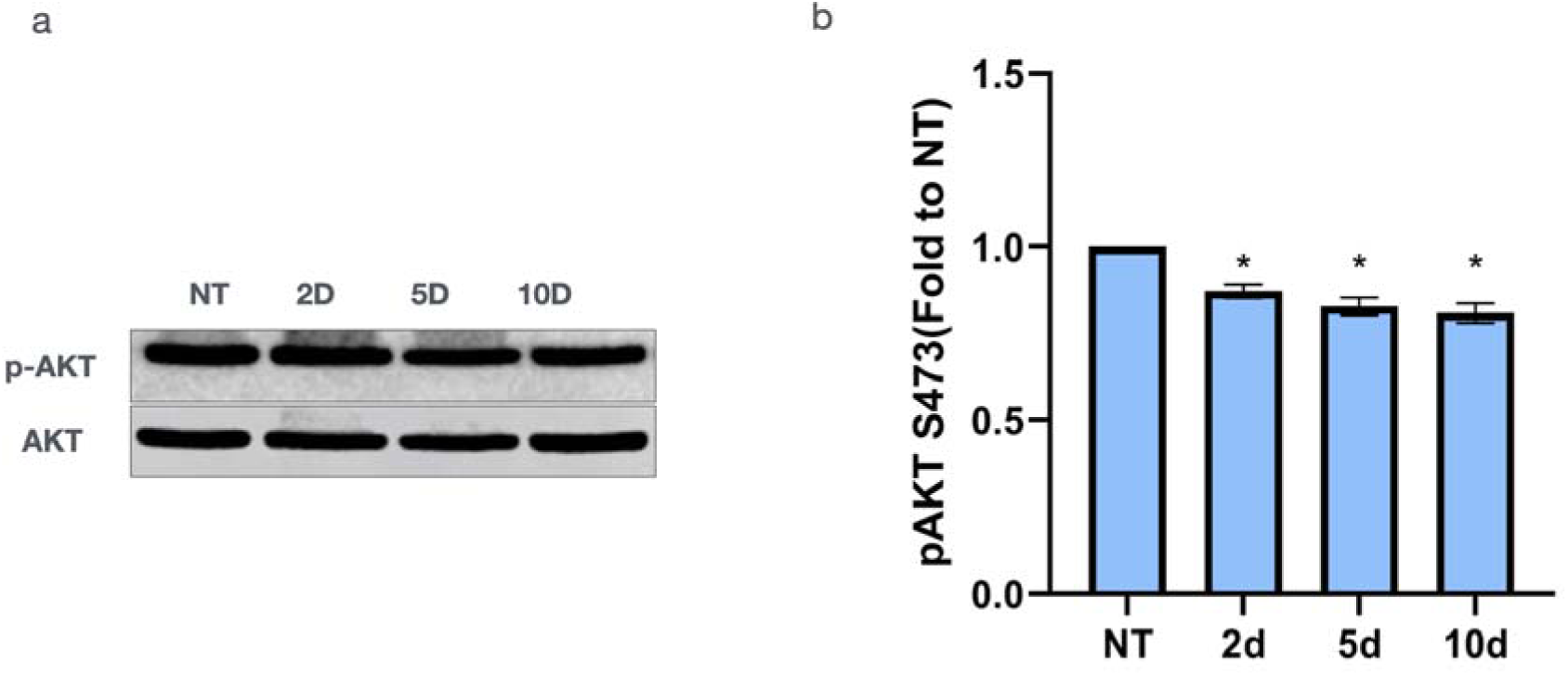
Rapamycin suppresses mTORC2 activity without altering mTORC1 signaling in MSCs. (a) Western blot membrane showing total AKT and p-AKT expression in MSCs treated with Rapamycin (10ng/ml) for 2, 5, and 10 days. (b) Quantification of p-AKT levels relative to total AKT. Rapamycin caused a time-dependent decrease in p-AKT expression. Data are represented as mean ± SEM bars. N = 10, \**P* < 0.05, compared to NT.

#### 3.7.2 mTORC2 inhibition modulated α-SMA and transgelin expression

Since rapamycin significantly reduced mTORC2 activity, we were interested in determining whether mTORC2 inhibition could increase the expression of α-SMA and transgelin in MSCs, as observed with rapamycin treatment. To test this, we selectively inhibited mTORC2 using JR-AB2-011, which blocks Rictor–mTOR interaction and reduces AKT Ser473 phosphorylation (Figure 6a). MSCs were treated with 25 μM JR-AB2-011 for 5 days, and the protein expression of α-SMA and transgelin determined. Inhibition of mTORC2 with JR-AB2-011 resulted in a 69% reduction in α-SMA expression and a 76% reduction of transgelin expression compared to control levels (*p < 0.05*) (Figure 6a-c). JR-AB2-011 treatment did not alter phospho-S6 (S235/236) levels, an indirect target of mTORC1 activation, but did reduce phosphor-AKT S473 levels. These results suggest that rapamycin-induced changes in α-SMA and transgelin expression are dependent on mTORC1 inhibition, rather than mTORC2.

**Figure 6:**
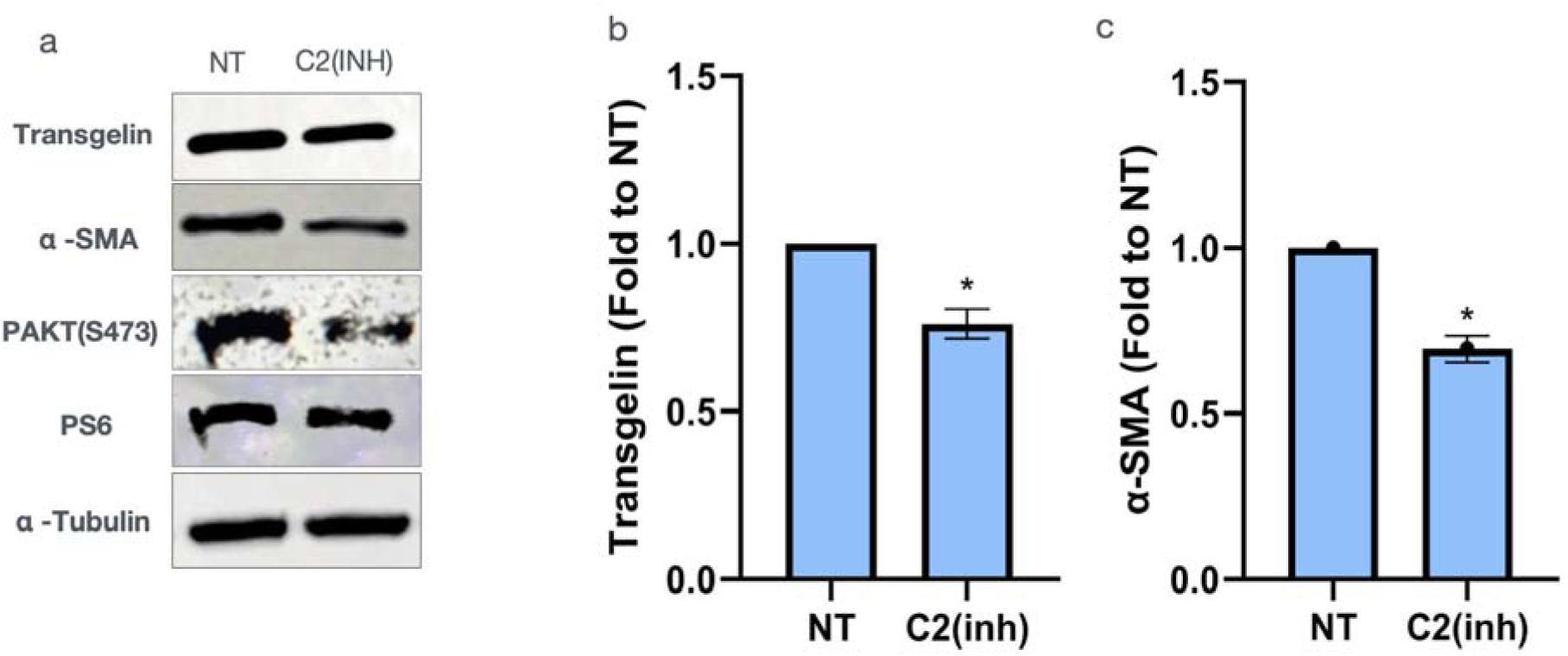
Selective mTORC2 inhibition reduces VSMC marker expression. (a) Western blot membrane showing α-SMA, transgelin, p-AKT(S473), PS6, and α-tubulin expression in MSCs treated with a mTORC2-specific inhibitor or no treatment group. (b) Quantification of α-SMA expression, showing a significant decrease (0.69-fold, p < 0.01) with mTORC2 inhibition. (c) Quantification of transgelin expression, showing a 0.76-fold decrease with mTORC2 inhibition. Data are represented as mean ± SEM bars. N = 10, \**P* < 0.05, compared to NT.

### 3.8 Rapamycin Induces Sproutations and Increases VEGFA Expression in MSCs

During rapamycin treatment, the sprouting from one cell to another was consistently observed in MSC cultures (Figure 7a). To further characterize this finding, we quantified the number of these sproutations observed in control and rapamycin-treated cultures. MSCs were treated with rapamycin or vehicle for 10 days and then labeled with Calcein AM to enhance visualization. The frequency was determined by randomly selecting 10 non-overlapping microscopic fields per dish from five independent biological replicates and counting the number of sproutations observed. As shown in Figure 7a and 7b, vehicle-treated control cells only formed few sproutations. In contrast, cell sprotuations in rapamycin-treated cells were frequently observed (Figure 7a), with a 12-fold increase in rapamycin-treated cells compared to controls (p < 0.05) (Figure 7b).

**Figure 7:**
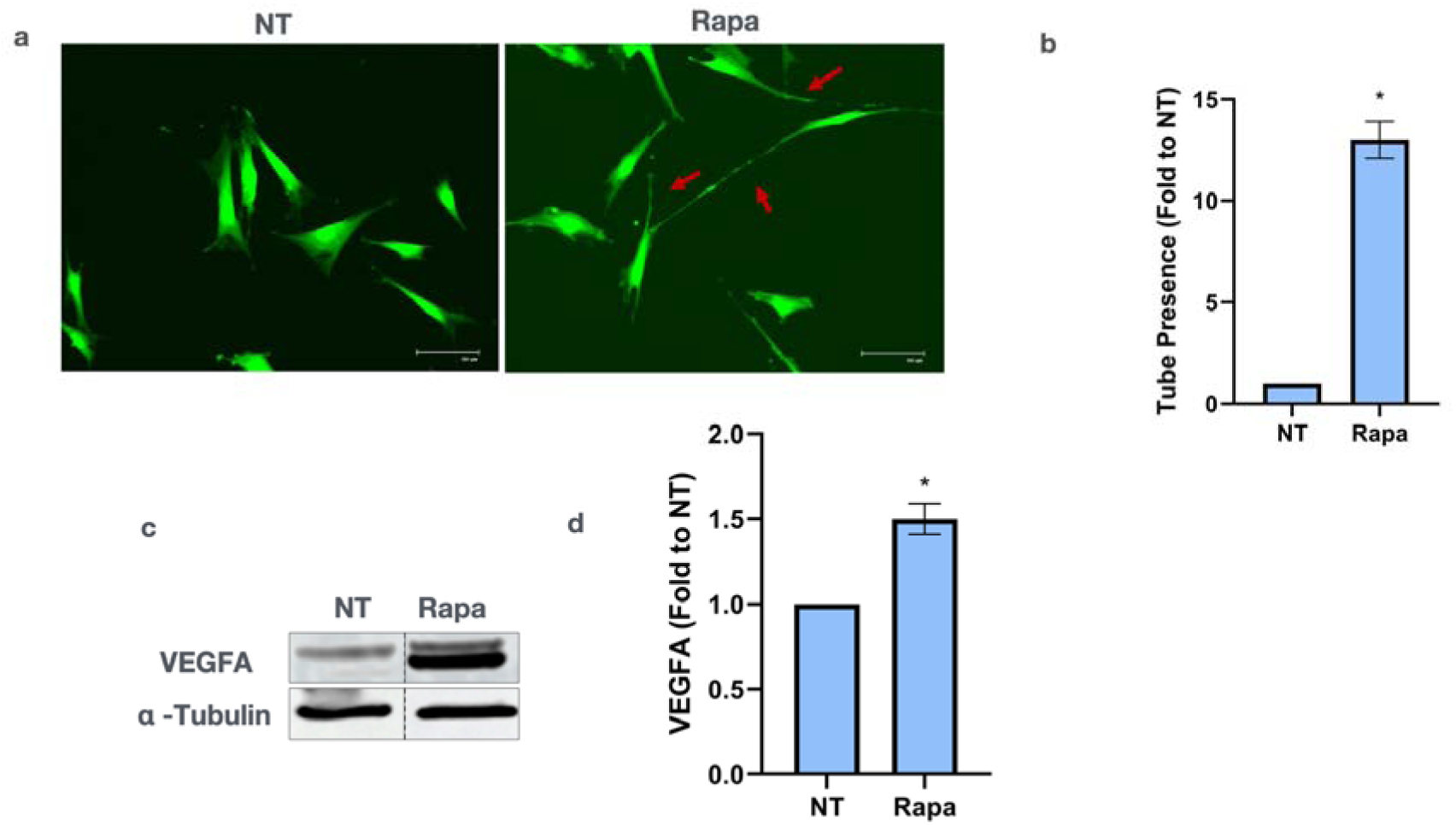
Rapamycin promotes projection structure formation and increases VEGFA expression in MSCs. (a) Fluorescence microscopy showing projections from MSCs treated with rapamycin compared to control. Green fluorescence (Calcein AM) highlights viable cells showing projections in MSCs treated with rapamycin (pointed by arrows). (b) Quantification of projection length showed a 13-fold increase in the rapamycin group compared to the control. (c) VEGFa protein expression in rapamycin group compared to no treatment showed a significant upregulation with 1.5-fold increase. Data are represented as mean ± SEM bars. N = 10, \**P* < 0.05, compared to NT.

Since rapamycin treatment induced cellular sprouting, we were interested in determining whether rapamycin induced endothelial and angiogenic gene expression in MSCs. Therefore, VEGFA protein expression was assessed following 10 days of rapamycin treatment at 10 ng/ml. As shown in Figure 7c and d, VEGFA expression was significantly increased 2.8-fold relative to controls (p < 0.05) (Figure 7c). Since VEGFA was upregulated, we further evaluated whether additional endothelial markers were induced. Therefore, western blot analysis was used to determine the expression of CD31, CD34, von Willebrand factor (VWF), and E-cadherin in control and rapamycin-treated cultures. However, these EC proteins were not detected in either control or rapamycin-treated groups (data not shown). Positive EC control lysates were used to confirm antibody performance (data not shown).

These results demonstrate that rapamycin induces the expression of the angiogenic factor VEGFA but not endothelial cell markers.

## 4. DISCUSSION

This study investigated the potential of rapamycin to increase MSCs expression of the VSMC proteins α-SMA and transgelin. We found that rapamycin reduced MSC proliferation but was not cytotoxic, even at high drug concentrations and with prolonged exposure. Rapamycin induced a profound increase in cell size, with cells adopting a more cuboidal morphology. Rapamycin treatment increased the expression of α-SMA and transgelin in MSCs and was required to maintain the expression of these genes. While rapamycin did not induce the expression of EC proteins, it did promote the expression of VEGFA, a potent angiogenic factor. Interestingly, rapamycin also induced the expression of sproutations from cells and connections between two neighboring cells. To our knowledge, this is the first study to demonstrate the potential of rapamycin to increase expression of VEGFA and VSMC contractile proteins α-SMA and transgelin in MSCs.

### 4.1 Proliferation and viability of MSCs under rapamycin

Rapamycin is a known inhibitor of cell proliferation (45–47) and as such has been successfully used in the treatment of restenosis in cardiac and peripheral artery diseases (48). Consistent with this, our results demonstrate that rapamycin treatment significantly reduced MSC proliferation rate. Our data also demonstrates that rapamycin was not cytotoxic for at least 1 week. Previous studies have shown that rapamycin is antiproliferative, inhibiting the progression from the G1 phase to the S phase, but not toxic to cells (49–51).

### 4.2 Rapamycin induced morphological changes and expression of VSMC genes in MSCs

Our results demonstrate that MSCs express VSMC genes, α-SMA, and transgelin under normal conditions and that rapamycin treatment upregulates the expression of these genes. However, the expression of α-SMA, and transgelin decreased after rapamycin withdrawal suggesting that rapamycin altered gene expression but did not induce MSC differentiation. This transient effect may be due to the reversal of mTORC1 blockade upon rapamycin withdrawal (52–54). Previous studies demonstrated that MSCs express VSMC genes, α-SMA, SM22alpha, calponin, and SM myosin heavy chain under the influence of transforming growth factor-β1(TGF-β1) (55, 56). Likewise, Wang et al. showed that adipose-derived stem cells can differentiate into VSMC when treated with TGF-β1 and bone morphogenic protein-4 (BMP-4) (56). Our results demonstrate an increase in cell area and a change in shape from an elongated spindle to a cuboidal form under rapamycin treatment.

### 4.3 Role of mTORC1 in modulating MSCs expression of VSMC protein expression

Rapamycin is a specific mTORC1 inhibitor which can indirectly inhibit mTORC2 activity during prolonged rapamycin treatment (35). To confirm that VSMC protein expression was being regulated by mTORC1, we determined activity of mTORC1 via changes in phospho-S6 ribosomal protein (p-S6) and Phospho-AKT (p-AKT). It is well established that pS6 is a downstream molecule and a good indicator of mTORC1 activity (57). Our results showed complete inhibition of pS6 in the treatment groups compared to the no-treatment groups, signifying that mTORC1 activity is entirely inhibited by rapamycin. To examine the effect of rapamycin on mTORC2, we checked the expression of p-AKT, whose activity is regulated by mTORC2 (58). Study results showed decreased expression of p-AKT, indicating the inhibition of mTORC2. When mTORC2 activity was blocked using a complex 2 specific antagonist, which blocks Rictor–mTOR interaction and reduces AKT Ser473 phosphorylation, it demonstrated a decreased expression of VSMC protein expression. This observed trend infers the role of mTORC1 in inducing morphological changes in MSCs and expressing VSMC protein expression. Complete inhibition of mTORC1 under rapamycin induces differentiation of MSCs, and the activity of mTORC1 has a negative effect on this differentiation.

Previous studies on rapamycin and MSC demonstrated the role of PI3K/Akt/mTOR signaling in phenotypic modulation of MSC to express VSMC marker expression, including L-type Ca(2+) channels in MSC, by inhibiting mTOR using rapamycin (51). To the best of our knowledge, this is the first study to specifically demonstrate the role of mTORC1 in the expression of VSMC contractile protein marker expression in MSCs. These findings open a new avenue for research exploring the therapeutic potential of MSCs in cerebrovascular disorders.

### 4.4 Rapamycin induces VEGFA expression in MSCs

Our results also indicated that rapamycin enhanced VEFGA expression. VEGFA plays a critical role in angiogenesis along with its receptor, vascular endothelial growth factor receptor (VEGFR) (59). VEGFA interacts with VEGFR2 to induce endothelial cell proliferation, migration, and survival, ultimately leading to angiogenesis (60, 61). A study by Beckermann et al. on pancreatic cancer demonstrated increased MSC-VEGF production which migrated to the cancer tissue after being injected into mice to promote angiogenesis in the cancerous tissue (62). Our VEGFA expression is also complemented by the change in the shape of MSCs under rapamycin. Staining of MSCs treated with rapamycin shows sprouting of MSC and the connection of two cells. Previous *in-vitro* studies on endothelial cells demonstrated increased sprouting between cells supporting the angiogenetic potential of MSCs (63) (64). Xu et al. showed that MSCs form tube-like structures and are involved in angiogenesis when treated with VEGF (65). These results denote rapamycin may potentially play a crucial role in angiogenesis by enhancing VEGFA expression in MSCs.

## 5. CONCLUSION

Several previous studies demonstrated the use of MSCs for the treatment of aneurysms (66–69). Liu et al. showed decreased ICA rupture rates in mice when micro vesicles derived from Human MSCs are injected into vasculature (66). The same effect of MSCs in preventing rupture of aneurysms was also demonstrated by Kuwabara and his team after intravenously injecting MSCs (70). Rouchaud and group showed improved saccular aneurysm healing rates in rabbit when treated with coils coated with adipose derived MSCs (68). These studies suggest the potential use of MSCs in the treatment of ICA.

The ultimate goal of ICA treatment is the reformation of the parent artery across the aneurysm neck, thereby removing the aneurysm from the circulation. Therefore, the ability of rapamycin to increase VEGFA expression in MSCs could allow for a novel MSC-based therapeutic intervention in cerebral aneurysm management. In this approach, EC recruitment, proliferation and survival are enhanced by rapamycin induced MSC VEGFA expression, thereby increasing cellular migration and growth across the aneurysm neck. Future studies should further investigate the therapeutic potential of rapamycin treated MSC in the treatment of ICA.

